# Coding and regulatory variants affect serum protein levels and common disease

**DOI:** 10.1101/2020.05.06.080440

**Authors:** Valur Emilsson, Valborg Gudmundsdottir, Alexander Gudjonsson, Mohd A Karim, Marjan Ilkov, James R. Staley, Elias F. Gudmundsson, Brynjolfur G. Jonsson, Lenore J. Launer, Jan H. Lindeman, Nicholas M. Morton, Thor Aspelund, John R. Lamb, Lori L. Jennings, Vilmundur Gudnason

**Affiliations:** Icelandic Heart Association, Holtasmari 1, IS-201 Kopavogur, Iceland; Faculty of Medicine, University of Iceland, 101 Reykjavik, Iceland; Wellcome Trust Sanger Institute, Welcome Genome Campus, Hinxton, Cambridgeshire CB10 1SA, UK; Open Targets, Wellcome Genome Campus, Hinxton, Cambridgeshire CB10 1SD, UK; MRC/BHF Cardiovascular Epidemiology Unit, Department of Public Health and Primary Care, University of Cambridge, Cambridge, UK; Laboratory of Epidemiology and Population Sciences, Intramural Research Program, National Institute on Aging, Bethesda, MD 20892-9205, USA; Department of General Surgery Leiden University Medical Center, Leiden. Holland; Centre for Cardiovascular Sciences, Queen’s Medical Research Institute, University of Edinburgh, Edinburgh EH16 4TJ, UK; GNF Novartis, 10675 John Jay Hopkins Drive, San Diego, CA 92121, USA; Novartis Institutes for Biomedical Research, 22 Windsor Street, Cambridge, MA 02139, USA

## Abstract

Circulating proteins are prognostic for human outcomes including cancer, heart failure, brain trauma and brain amyloid plaque burden. A deep serum proteome survey recently revealed close associations of serum protein networks and common diseases. The present study reveals unprecedented number of individual serum proteins that overlap genetic signatures of diseases emanating from different tissues of the body. Here, 54,469 low-frequency and common exome-array variants were compared with 4782 protein measurements in the serum of 5343 individuals of the deeply annotated AGES Reykjavik cohort. Using a study-wide significant threshold, 2019 independent exome array variants affecting levels of 2135 serum proteins were identified. These variants overlapped genetic loci for hundreds of complex disease traits, emphasizing the emerging role for serum proteins as biomarkers of and potential causative agents of multiple diseases.

---

Large-scale genome-wide association studies (GWASs) have expanded our knowledge of the genetic basis of complex disease. As of 2018, approximately 5687 GWASs have been published revealing 71,673 DNA variants to phenotype associations^1^. Furthermore, exome-wide genotyping arrays have linked rare and common variants to many complex traits. For example, 444 independent risk variants were recently identified for lipoprotein fractions across 250 genes^2^. Despite the overall success of GWAS, the common lead SNPs rarely point directly to a clear causative polymorphism, making determination of the underlying disease mechanism difficult^3–6^. Regulatory variants affecting mRNA and/or protein levels and structural variants like missense mutations can point directly to the causal candidate. Alteration of the amino acid sequence may affect protein activity and/or influence transcription, translation, stability, processing, and secretion of the protein in question^7–9^. Thus, by integrating intermediate traits like mRNA and/or protein levels with genetics and disease traits, the identification of the causal candidates can be enhanced^3–6^.

Proteins are arguably the ultimate players in all life processes in disease and health, however, high throughput detection and quantification of proteins has been hampered by the limitations of available proteomic technologies. Recently, a custom-designed Slow-Off rate Modified Aptamer (SOMAmer) protein profiling platform was developed to measure 4782 proteins encoded by 4137 human genes in the serum of 5457 individuals from the AGES Reykjavik study (AGES-RS)^10^, resulting in 26.1 million individual protein measurements. Various metrics related to the performance of the proteomic platform including aptamer specificity, assay variability and reproducibility have already been described^10^. We demonstrated that the human serum proteome is under strong genetic control^10^, in line with findings of others applying identical or different proteomics technologies^11,12^. Moreover, serum proteins were found to exist in regulatory groups of network modules composed of members synthesized in all tissues of the body, suggesting that system level coordination or homeostasis is mediated to a significant degree by thousands of proteins in blood^13^. Importantly, the deep serum and plasma proteome is associated with and prognostic for various diseases as well as human life span^10,14–20^.

Here, we regressed levels of 4782 proteins on 54,469 low-frequency and common variants from the HumanExome BeadChip exome array, in sera from 5343 individuals of the deeply phenotyped AGES-RS cohort. Further cross-referencing of all significant genotype-to-protein associations to hundreds of genetic loci for various disease endpoints and clinical traits, demonstrated profound overlap between the genetics of circulating proteins and disease related phenotypes. We highlight how triangulation of data from different sources can link genetics, protein levels and disease(s), with the intention of cross-validating one another and point to potentially causal relationship between proteins and complex disease(s).

Using genotype data from an exome array (HumanExome BeadChip) enriched for structural variants and tagged for many GWAS risk loci (**Methods**), the effect of low-frequency and common variants on the deep serum proteome was examined. Quality control filters^21^, and exclusion of monomorphic variants reduced the available variants to 76,891. Additionally, we excluded variants at minor allele frequency (MAF) < 0.001 as they provide insufficient power for single-point association analysis^22^. This resulted in 54,469 low-frequency (54%, MAF<0.05) and common variants (46%, MAF ≥0.05) that were tested for association to each of the 4782 human serum protein measurements using linear regression analysis adjusted for the confounders age and sex (**Methods**). The current platform targets the serum proteome arising largely from active or passive secretion, ectodomain shedding, lysis and/or cell death^10,23^. **Figure 1a** highlights the classification of the protein population targeted by the aptamer-based profiling platform, showing over 68.7% of the proteins are secreted or single pass transmembrane (SPTM) receptors.

**Fig. 1.**
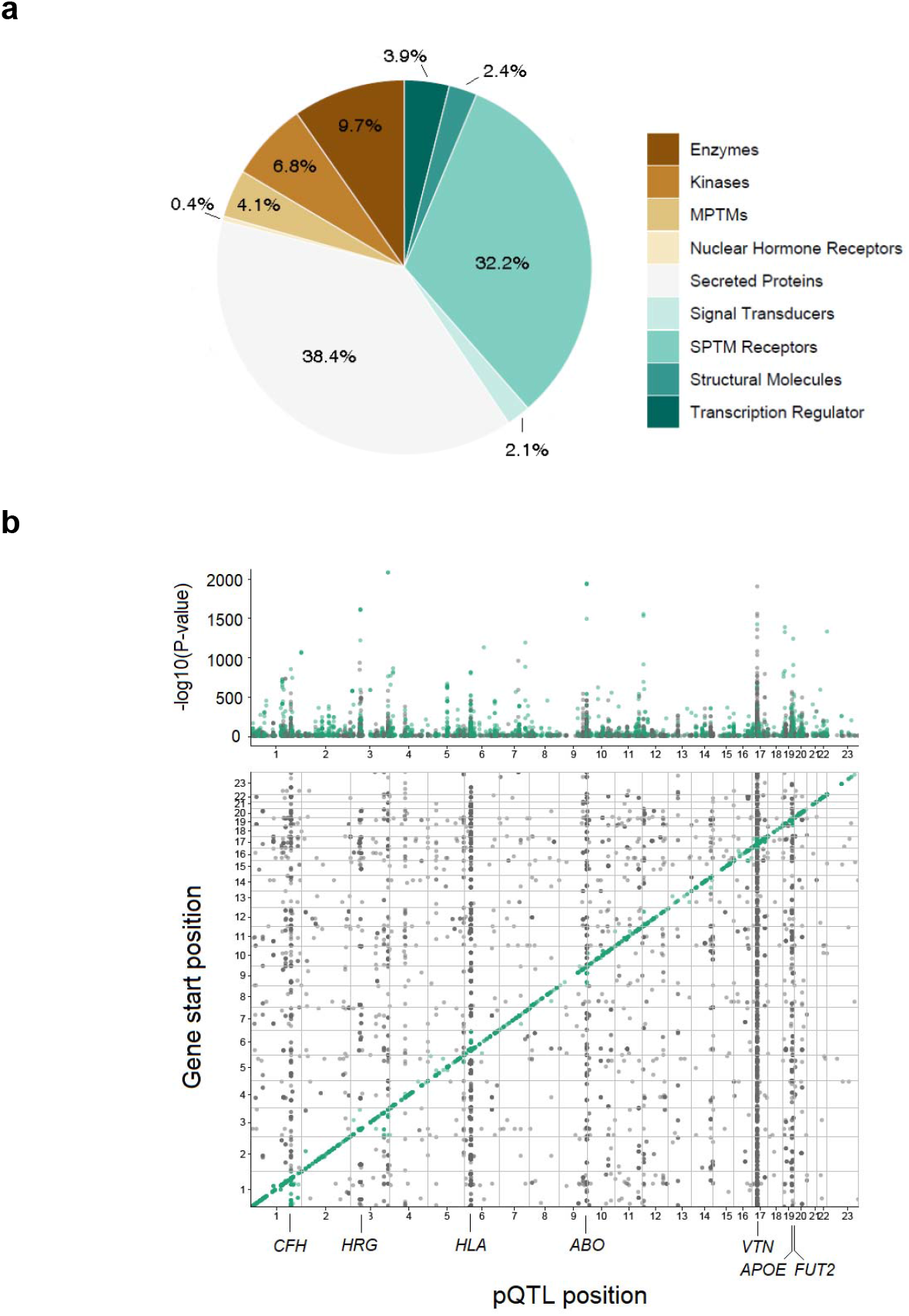
Classification of the target protein population and genomic locations of observed pQTLs. **a**. Pie chart showing the relative distribution (percentage) of the different protein classes targeted by the present proteomics platform, with secreted proteins (38.4%) and single pass transmembrane (SPTM) receptors (32.2%) dominating the target protein population. Protein classes were manually curated based on information from the SecTrans, Gene Ontology (GO) and Swiss-Prot databases, and were composed of secreted proteins (e.g. cytokines, adipokines, hormones, chemokines and growth factors), SPTM receptors (e.g. tyrosine and serine/threonine kinase receptors), multi-pass transmembrane (MPTM) receptors (e.g. GPCR, ion channels, transporters), enzymes (intracellular), kinases, nuclear hormone receptors, structural molecules, transcriptional regulators and signal transducers. **b**. The Manhattan plot in the top panel uses precise P-values to highlight all study-wide significant associations in Supplementary Table S1. The bottom panel shows the genomic locations of all study-wide significant pQTLs (P < 1.92×10^-10^), where the start position of the protein encoding gene is shown on the y-axis and the location of the pSNP at the x-axis. *Cis* acting effects, using a 300kb window, appear at the diagonal while *trans* acting pQTL effects including *trans* hot spots show up off-diagonally. The genetic loci highlighted across the x-axis are *trans*-acting hotspots.

Applying a Bonferroni corrected significance threshold of P < 1.92×10^-10^ (0.05/54469/4782) we detected 5472 exome array variants that were associated with variable levels of 2135 serum proteins (**Supplementary Table 1** and **Fig. 1b**), of which 2019 variants are independent (**Supplementary Table S2**). **Supplementary Table 1** lists all associations at P-value < 1×10^-6^, or 10,200 exome array variants affecting 3096 human proteins. These protein quantitative trait loci (pQTLs) were *cis* and/or *trans* acting including several *trans* acting hotspots with pleiotropic effects on multiple co-regulated proteins (**Fig. 1b**). Secreted proteins were enriched for pQTLs (P-value < 0.0001) as compared to non-secreted proteins using 10,000 permutations to obtain the empirical distribution of the χ^2^ test of equality of proportions (**Supplementary Fig. S1**). This suggests that proteins bound for the systemic environment are subject to more genetic regulation than other proteins identified by the current platform. **Supplementary Table S3** summarizes various pathogenicity prediction scores for all independent study-wide significant pQTLs in **Supplementary Table S2**, using the Ensembl Variant Effect Predictor (VEP)^24,25^. Next, we cross-referenced all the 5472 study-wide significant pQTLs with a comprehensive collection of genetic loci associated with diseases and clinical traits from the curated PhenoScanner database^26^, revealing that 60% of all pQTLs were linked to at least one disease-related trait (**Supplementary Table S4**). We have shown in our previous studies that genetic loci affecting several serum proteins exhibit pleiotropy in relation to complex diseases^10^. An example of a possible pleiotropic effect mediated by the variant rs2251219 within the gene *PBRM1* affecting multiple proteins and sharing genetics with various diseases and clinical features is illustrated in **Fig. 2**. **Supplementary Fig. S2** depicts the relationship between all proteins and some quantitative traits associated with rs2251219. **Table 1** highlights a selected set of pQTLs that share genetics with diseases of different etiologies including disorders of the brain, metabolism, immune and cardiovascular system and cancer. In the sections that follow, we give examples of serum pQTLs that overlap disease risk loci and demonstrate how different data sources can cross-validate one another. Although data triangulation can be used to infer directional consistency, it cannot tell whether the relationship is causal or reactive to a given outcome. As a result, we used two-sample Mendelian randomization analysis (MR) on highlighted examples to test support for a protein’s causality to an outcome.

**Fig. 2.**
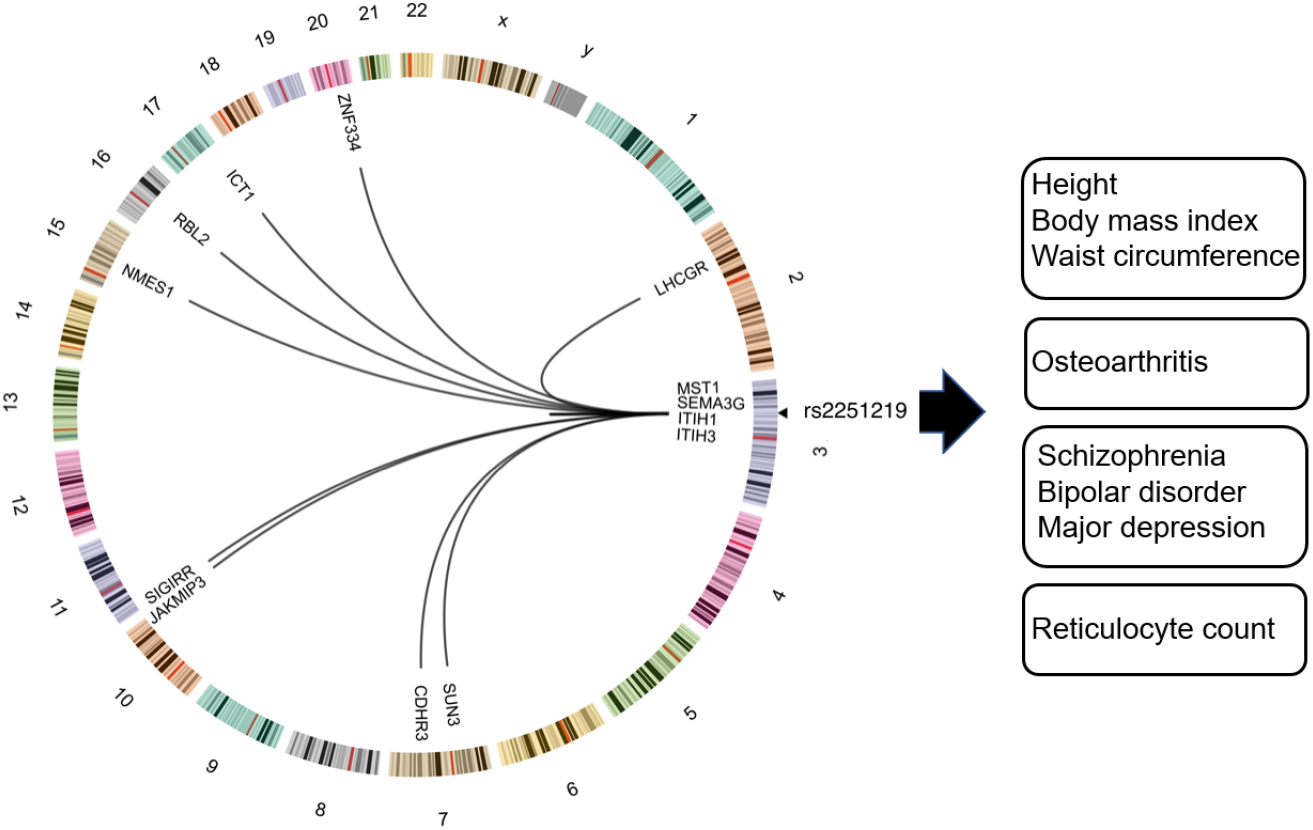
Pleiotropy of rs2251219 affecting many proteins and disease traits. **a**. Circos plot showing the effect of the variant rs2251219 (Supplementary Tables 1 and 2) on 13 proteins acting in *cis* or *trans* and sharing genetics with various diseases of different etiologies. Only study-wide significant (P < 1.92 x 10^-10^) genotype-to-protein associations are shown. Lines going from rs225121show links to genomic locations of the protein encoding genes affected while numbers refer to chromosomes.

**Table 1.** Selected examples of exome array variants affecting serum protein levels and complex disease. CHD, coronary heart disease; VTE, venous thromboembolism; CKD, chronic kidney disease; T2D, type 2 diabetes; VAT, visceral adipose tissue; LOAD, late-onset Alzheimer’s disease; SLE, systemic lupus erythematous; IBD, inflammatory bowel disease; AMD, age-related macular degeneration; N/A, not applicable. All reported effects are genome-wide significant at P < 1.92×10^-10^.

| Disease class | Disease trait | PMID or database | pQTL | GWAS lead SNP(s)<sup>a</sup> | Function pSNP<sup>b</sup> | Mapped GWAS locus<sup>c</sup> | #Proteins affected | Example of *cis* and/or *trans* affected proteins<sup>d</sup> |
| --- | --- | --- | --- | --- | --- | --- | --- | --- |
| *Cardiovascular* |  |  |  |  |  |  |  |  |
|  | CHD | 28714975 | rs12740374 | rs12740374 | 3'-UTR | CELSR2 | 8 | **C1QTNF1, IGFBP1** |
|  | VTE | UKBB, 28373160 | rs2343596 | rs16873402, rs4602861 | Intron | ZFPM2 | 7 | **VEGFA, DKK1** |
|  | Stroke | 26708676 | rs653178 | rs653178 | Intron | ATXN2 | 2 | **THPO, CXCL11** |
| *Metabolic* |  |  |  |  |  |  |  |  |
|  | T2D | 22885922 | rs7202877 | rs7202877 | Intergenic | CTRB1 | 5 | **CTRB1, PRSS2, CPB1** |
|  | VAT | 20935629 | rs9491696 | rs9491696 | Intron | RSPO3 | 1 | RSPO3 |
|  | Triglyceride | 21386085 | rs2266788 | rs2266788 | 3'-UTR | APOA5 | 5 | **APOA5, PCSK7, ANGPTL3** |
| *CNS* |  |  |  |  |  |  |  |  |
|  | LOAD | 21460840 | rs610932 | rs610932 | 3'-UTR | MS4A6A | 3 | **TREM2, GLTPD2** |
|  | Parkinson | 21738487 | rs6599389 | rs6599389 | Intron | GAK | 1 | IDUA |
|  | Schizophrenia | 25056061 | rs3617 | rs3617 | Q315K | ITIH3 | 8 | **ITIH3, JAKMIP3** |
| *Inflammatory* |  |  |  |  |  |  |  |  |
|  | SLE, T1D | 26502338 | rs2304256 | rs2304256 | V362F | TYK2 | 2 | ICAM1, ICAM5 |
|  | Crohn's, IBD | 21102463 | rs11209026 | rs11209026 | R381Q | IL23R | 1 | IL23R |
|  | AMD | 2355636 | rs10737680 | rs10737680 | Intron | CFH | 22 | **CFH, CFHR1, CFB** |
| *Cancer* |  |  |  |  |  |  |  |  |
|  | Colorectal | 24836286 | rs2241714 | rs1800469 | I11M | TMEM91 | 3 | **B3GNT2, TGFB1** |
|  | Lung | 18978787 | rs3117582 | rs3117582 | Intron | APOM | 10 | **MICB, ISG15** |
|  | Melanoma | 18488026 | rs910873 | rs910873 | Intron | PIGU | 1 | ASIP |

<sup>a</sup>Protein QTLs overlapping GWAS lead SNPs using the PhenoScanner database<sup>23</sup>. No SNP proxies were applied except when the lead pSNP was not in the query then we used the best proxy ( $r^2 \geq 0.8$ between markers). <sup>b</sup>The functional annotation of pQTLs was obtained from the PhenoScanner database<sup>23</sup>. <sup>c</sup>Reported causal candidates are from the GWAS Catalog<sup>67</sup>. <sup>d</sup>The definition of *cis* vs. *trans* effects is somewhat arbitrary depending on the window size chosen across the protein gene in question. However, all affected proteins located at other chromosomes than the pQTL location, were considered *trans* acting and are highlighted in bold letters. All significant pQTLs are listed in Supplementary Table 1 and the overlap with GWAS risk loci summarized in Supplementary Table 4.

Variable levels of the anti-inflammatory protein TREM2 were associated with two distinct genomic regions (**Fig. 3a** and **Supplementary Fig. S3**). This included the missense variant rs75932628 (NP_061838.1: p.R47H) in *TREM2* at chromosome 6 (**Fig. 3b**), known to confer a strong risk of late-onset Alzheimer’s disease (LOAD)^27^. The variant was also associated with IGFBPL1 (P = 3×10^-18^) in serum (**Supplementary Table 1**), a protein recently implicated in axonal growth^28^. Intriguingly, the region at chromosome 11 associated with soluble TREM2 levels harbors variants adjacent to the genes *MS4A4A* and *MS4A6A* including rs610932 known to influence genetic susceptibility for LOAD^29^ (**Table 1** and **Fig. 3a, b)**. The variant rs610932 was also associated with the proteins GLTPD2 and A4GALT (**Supplementary Table 1**). The alleles increasing risk of LOAD for both the common variant rs610932 and the low-frequency variant rs75932628 were associated with low levels of soluble TREM2 (**Fig. 3b**). Consistently, we find that the high-risk allele for rs75932628 was associated with accelerated mortality post incident LOAD in the AGES-RS (**Fig. 3c**). It is of note that the levels of TREM2 in the cerebrospinal fluid (CSF) reflect the activity of brain TREM2-triggered microglia^4,30^, while high levels of CSF TREM2 have been associated with improved cognitive functioning^31^. **Supplementary Fig. S4** highlights the correlation (Spearman rank) between the different proteins affected by the LOAD risk loci at chromosomes 6 and 11. The accumulated data show a directionally consistent effect at independent risk loci for LOAD converging on the same causal candidate TREM2. Furthermore, a two-sample MR analysis using genetic instruments across the *TREM2* and *MS4A4A/MS4A6A* loci and GWAS associations for LOAD in Europeans as outcome^32^, provided evidence that variable TREM2 protein levels are causally related to LOAD (P = 7.6×10^-5^) (**Fig 3d**). In summary, these results demonstrate that the effect of genetic drivers on major brain-linked disease like LOAD can be readily detected in serum to both inform on the causal relationship and the directionality of the risk mediating effect. This would also suggest that serum may be an accessible proxy for microglia function and cognition.

**Fig. 3.**
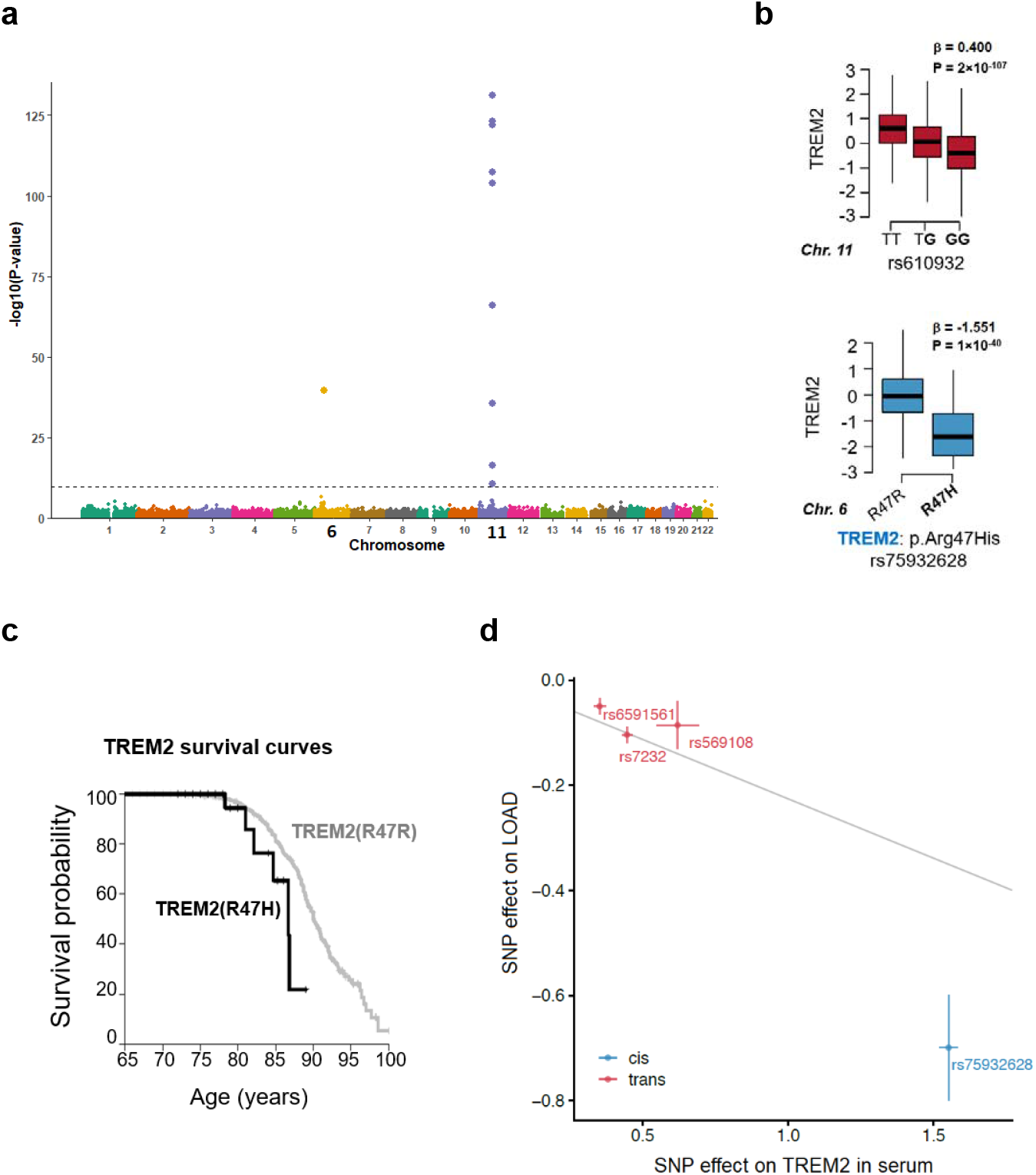
Effects of distinct risk loci for LOAD converge on the protein TREM2. **a.** The Manhattan plot highlights variants at two distinct chromosomes associated with serum TREM2 levels. Study-wide significant associations at P < 1.92×10^-10^ are indicated by the horizontal line. The y-axis shows the -(log_10_) of the P-values for the association of each genetic variant on the exome array present along the x-axis. Variants at both chromosomes 6 and 11 associated with TREM2 have been independently linked to risk of LOAD including the rs75932628 (NP_061838.1: p.R47H) in TREM2 at chromosome 6 and the variant rs610932 at chromosome 11. **b.** Boxplot of the *trans* effect of the well-established GWAS risk variant rs610932 for LOAD on TREM2 serum levels (upper panel), where the LOAD risk allele G (highlighted in bold) is associated with lower levels of TREM2. Similarly, the LOAD causing p.R47H mutation was associated with low levels of TREM2 (lower panel). **c**. TREM2p.R47H carriers demonstrated lower survival probability post-incident LOAD compared to TREM2p.R47R carriers (P = 0.04). **d**. Scatterplot for the TREM2 protein supported as having a causal effect on LOAD in a two sample MR analysis. The figure demonstrates the estimated effects (with 95% confidence intervals) of their respective *cis*- and *trans*-acting genetic instruments on the serum TREM2 levels in AGES-RS (x-axis) and risk of LOAD through a GWAS by Kunkle et al.^32^ (y-axis), using 21,982 LOAD cases and 41,944 controls. The line indicates the inverse variance weighted causal estimate (β = −0.226, SE = 0.057, P = 7.6×10^-5^).

Variable levels of the cell adhesion protein SVEP1 are associated with variants located at chromosomes 1 and 9 (**Supplementary Table 1**, **Fig. 4a** and **Supplementary Fig. S5**). Genetic associations to SVEP1 levels at chromosome 9 include the low-frequency missense variant rs111245230 in SVEP1 (NP_699197.3: pD2702G) (**Fig. 4b**), which was recently linked to coronary heart disease (CHD), blood pressure and type-2-diabetes (T2D)^33^. Overall, we found eight different missense mutations in *SVEP1* that were associated with SVEP1 serum levels (**Supplementary Table 1**). The CHD and T2D risk allele (C) of rs111245230 was associated with elevated levels of SVEP1, and SVEP1 levels were consistently elevated in CHD and T2D patients (**Fig. 4c**). Furthermore, high SVEP1 levels were positively correlated with systolic blood pressure (β = 2.10, P = 4×10^-12^) (**Fig. 4c**), but not with diastolic blood pressure (β = 0.115, P = 0.413). Consistently, higher serum levels of SVEP1 were associated with increased mortality post-incident CHD in the AGES-RS (HR = 1.27, P = 9×10^-9^) (**Fig. 4d**). The variants at chromosome 1 linked to SVEP1 levels (**Fig. 4a**), have not previously been linked to any disease. Given the currently available GWAS summary statistics, a two-sample MR analysis using *cis*-variants on chromosome 9 for SVEP1 as instruments and a GWAS associations for T2D^34^ support a causal relationship of SVEP1 with T2D (P = 1.2×10^-5^) (**Fig. 4e**), but not with CHD^35^ or systolic blood pressure^36^ (P > 0.05). Our data triangulation and causal tests integrating genetics, serum protein levels and disease(s), indicate that SVEP1 may be a therapeutic target for T2D.

**Fig. 4.**
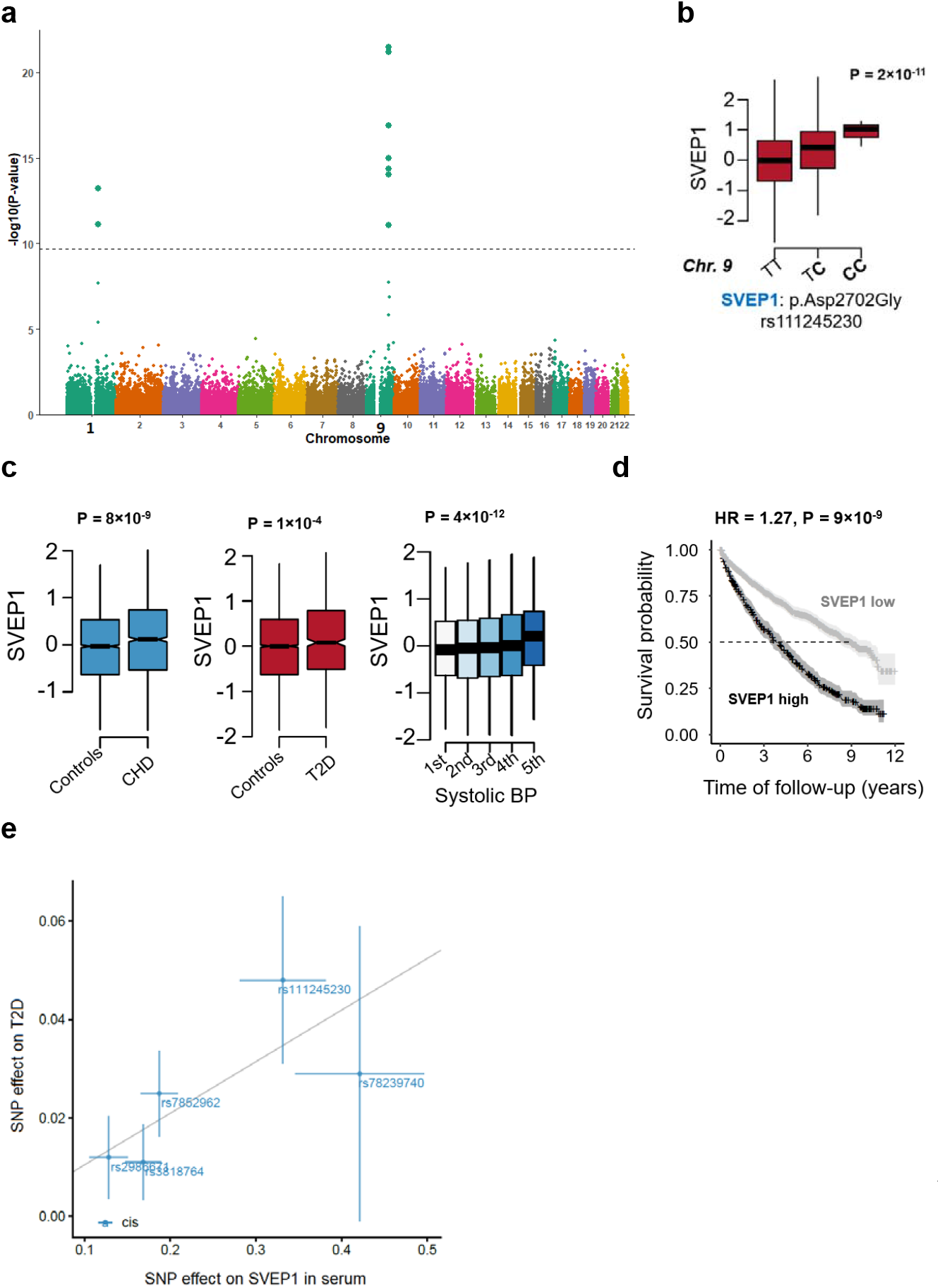
Variants affecting SVEP1 levels are associated with CHD and T2D. **a**. The Manhattan plot reveals variants at chromosomes 1 and 9 associated with serum SVEP1 levels. Study-wide significant associations at P < 1.92×10^-10^ are indicated by the horizontal line. The y-axis shows the - (log_10_) of the P-values for the association of each genetic variant on the exome array present along the x-axis. **b**. One of the variants associated with SVEP1 levels and underlying the peak at chromosome 9 is the low-frequency CHD risk variant rs111245230 (NP_699197.3: pAsp2702Gly). The CHD risk allele C (highlighted in bold) is associated with increased serum SVEP1 levels. **c**. Serum levels of SVEP1 were associated with CHD (P = 8×10^-9^), T2D (P = 1×10^-4^) and systolic blood pressure (P = 4×10^-12^) in the AGES-RS, all in a directionally consistent manner. **d**. Consistent with the directionality of the effects described above, we find that elevated levels of SVEP1 were associated with higher rates of mortality post-incident CHD. **e**. Scatterplot for the SVEP1 protein supported as having a causal effect on T2D in a two-sample MR analysis. The figure demonstrates the estimated effects (with 95% confidence intervals) of the SNP effect on serum SVEP1 levels and T2D from a GWAS in Europeans^34^ (y-axis), with 74,124 T2D patients and 824,006 controls. The line indicates the inverse variance weighted causal estimate (β = 0.105, SE = 0.024, P = 1.2×10^-5^).

The ILMN exome array contains several tags related to previous GWAS findings^37^, including many risk loci for cancer. For example, 21 loci associated with melanoma^38^ and 50 loci associated with colorectal cancer^39^. The exome array variant rs910873 located in an intron of the GPI transamidase gene *PIGU* was previously linked to melanoma risk^40^. The reported candidate gene *PIGU* is the gene most proximal to the lead SNP rs910873 and may be a novel candidate gene involved in melanoma. However, a more biologically relevant candidate is the agouti-signaling protein (*ASIP*) gene that is located 314kb downstream of the lead SNP rs910873. *ASIP* is a competitive inhibitor of MC1R^41^, and is thus strongly biologically implicated in melanoma risk^42^. We found that the melanoma risk allele for rs910873 was associated with elevated ASIP serum levels (P = 3×10^-179^) and the variant had no effect on other proteins measured with the current proteomic platform (**Fig. 5a**, **Supplementary Table 1** and **Table 1**). Interestingly, the pQTL rs910873 is also an eQTL for *ASIP* gene expression in skin^43^, showing directionally consistent effect on the mRNA and protein. Importantly, we found that serum ASIP levels were supported as causally related to malignant melanoma (P = 4.8 x 10^-26^) using a two-sample MR analysis on the protein-to-outcome causal sequence of events (**Fig. 5b**). Our data point to the ASIP protein underlying the risk at rs910873, thus providing supportive evidence for the hypothesis that ASIP mediated inhibition of MC1R results in suppression of melanogenesis and increased risk of melanoma^44^. An additional example is the susceptibility variant rs1800469 for colorectal cancer^45^, which is a proxy to the pQTL rs2241714 (*r*^2^=0.978) (**Table 1** and **Fig. 5b**). While the *TMEM91* gene was the reported candidate gene for the colorectal cancer risk at the rs1800469 (**Table 1**), we find that the risk variant affected three proteins in either *cis* (B3GNT8 and TGFB1) or *trans* (B3GNT2) (**Fig. 5b**). Intriguingly, all three proteins have previously been implicated in colorectal cancer^46–48^. Due to a lack of available and powered GWAS summary statistics data, we were unable to formally test the causality of these proteins to colorectal cancer. In conclusion, while we cannot rule out *PIGU* as a candidate gene for malignant melanoma, these findings point to an alternate, and possibly more biologically relevant, candidate, ASIP.

**Figure 5.**
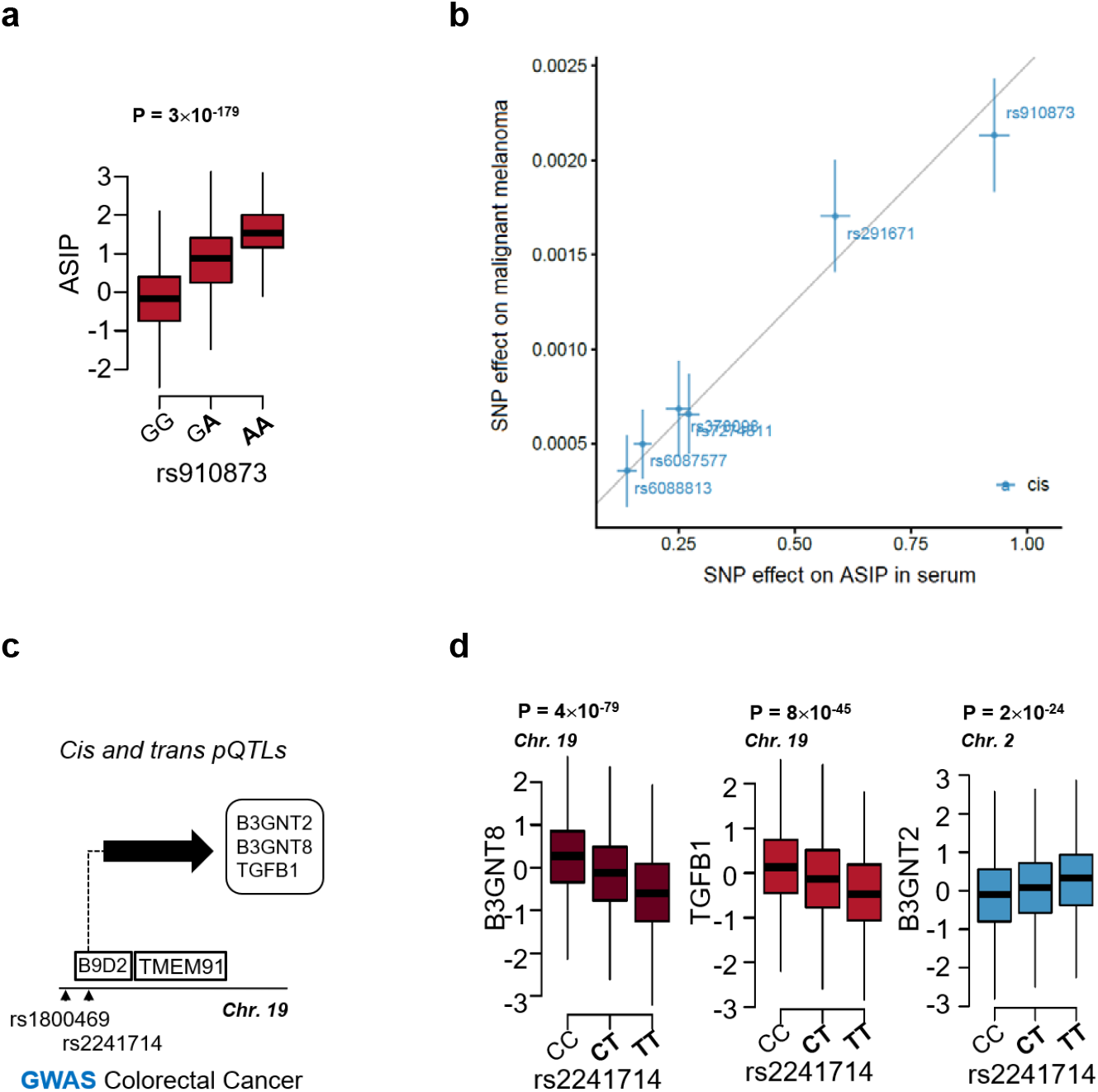
Proteins associated with malignant melanoma and colorectal cancer. **a**. The melanoma risk allele A (highlighted in bold) for the variant rs910873 is associated with high serum levels of ASIP. **b**. Scatterplot for the ASIP protein supported as having a causal effect on malignant melanoma in a two sample MR analysis. The figure demonstrates the estimated effects (with 95% confidence intervals) of their respective genetic instruments on the serum ASIP levels in AGES (x-axis) and risk of melanoma in GWAS by UK biobank data (UKB-b-12915) ^65^ (y-axis), that included 3598 melanoma cases and 459,335 controls. The line indicates the inverse variance weighted causal estimate (β = 0.0025, SE = 0.0002, P = 4.8×10^-26^). **c**. The pQTL rs2241714 is a proxy for the colorectal cancer associated variant rs1800469 (*r*^2^ = 0.978) (Supplementary Table 2), located within the gene *B9D2* and proximal to *TMEM91* which is the reported candidate gene at this locus (see Table 1). **d**. The variant rs2241714 (and rs1800469) regulate three serum proteins, B3GNT2 (in *trans*), B3GNT8 (in *cis*) and TGFB1 (in *cis*).

We outlined the construction of the serum protein network in our previous report and identified common genetic variants underlying the network structure^10^. This included a targeted study of the effects of common *cis* and *cis*-to-*trans* acting variants on levels of serum proteins. The comparison between that study and the current one using all independent study-wide signficiant associations (**Supplementary Table S2**) and linkage disequilibrium (LD) thresold of r^2^>0.50 for known associations, shows that 77.2% of the current study’s variant-to-protein associations are novel. Importantly, while 70% of the variants detected with the exome array are exonic and 59% of mapped pQTLs in the current study are exonic, only 7% of the identified pQTLs were exonic in our earlier report^10^. Previously, we discovered that 80% of *cis* pQTL effects and 74% of *trans* pQTL effects were replicated across populations and proteomics platforms measuring common variants^10^. Given that the exome array platform is enriched for rare and low-frequency variants, a comparable test of replication is not straightforward. Examining the proteins and variants measured across studies, we find that 76.0% of SNP-to-protein associations are novel in the present study when compared to, say, Sun et al.^11^, and 60.1% are novel when compared to the majority of studies published to date (**Supplementary Table S5**), for all independent associations in the current study and LD of *r*^2^<0.5 between study specific markers.

We report here that many of the measured serum proteins under genetic control share genetics with a variety of clinical features, including major diseases arising from various body tissues. This is in line with a recent population-scale survey of human induced pluripotent stem cells, demonstrating that pQTLs are 1.93-fold enriched in disease risk variants compared to a 1.36-fold enrichment for eQTLs^12^, underscoring the added value in pQTL mapping. We reaffirm widespread associations between genetic variants and their cognate proteins as well as distant *trans*-acting effects on serum proteins and demonstrate that many proteins are often involved in mediating the biological effect of a single causal variant affecting complex disease. Protein coding variants may cause technical artifacts in both affinity proteomics and mass spectrometry^49,50^. Systematic conditional and colocalization studies have shown, however, that pQTLs powered by common missense variants being artifactual are not a common event using the aptamer-based technology^11,51^, however, given the enrichment of missense variants in the present study, it may occur in some cases.

We note that with the ever-increasing availability of large-scale omics data aligned with the human genome, cross-referencing different datasets can result in findings that occurred by sheer chance. Hence, a systematic colocalization analysis has been proposed for identifying shared causal variants between intermediate traits and disease endpoints^52^. This is, however, not feasible for application of the exome array given its sparse genomic coverage. Instead, multi-omics data triangulation to infer consistency in directionality, the approach used in the present study, can enhance confidence in the causal call and offer insights and guidelines for experimental follow-up studies. In fact, the causal calls for TREM2 (LOAD), SVEP1 (T2D) and ASIP (melanoma) were validated, using a two-sample MR analysis. We previously asserted that serum proteins are intimately connected to and may mediate global homeostasis^10^. The accumulated data show that serum proteins are under strong genetic control and closely associated with diseases of different aetiologies, which in turn suggests that serum proteins may be significant mediators of systemic homeostasis in human health and disease.

## METHODS

### Study population

Participants aged 66 through 96 are from the Age, Gene/Environment Susceptibility Reykjavik Study (AGES-RS) cohort^53^. AGES-RS is a single-center prospective population-based study of deeply phenotyped subjects (5764, mean age 75±6 years) and survivors of the 40-year-long prospective Reykjavik study (n~18,000), an epidemiologic study aimed to understand aging in the context of gene/environment interaction by focusing on four biologic systems: vascular, neurocognitive (including sensory), musculoskeletal, and body composition/metabolism. Descriptive statistics of this cohort as well as detailed definition of the various disease endpoints and relevant phenotypes measured have been published^10,53^. The AGES-RS was approved by the NBC in Iceland (approval number VSN-00-063), and by the National Institute on Aging Intramural Institutional Review Board, and the Data Protection Authority in Iceland.

### Genotyping platform

Genotyping was conducted using the exome-wide genotyping array Illumina HumanExome-24 v1.1 Beadchip from Illumina (San Diego, CA, USA) for all AGES-RS participants as previously described^54^. The exome array was enriched for exonic variants selected from over 12,000 individual exome and whole-genome sequences from different study populations^37^, and includes as well tags for previously described GWAS hits, ancestry informative markers, mitochondrial SNPs and human leukocyte antigen tags^37^. A total of 244,883 variants were included on the exome array. Genotype call and quality control filters including call rate, heterozygosity, sex discordance and PCA outliers were performed as previously described^2,21^. Variants with call rate <90% or with Hardy–Weinberg P values < 1×10^−7^ were removed from the study. 72,766 variants were detected in at least one individual of the AGES-RS cohort. Of these variants, 54,469 had a minor allele frequency > 0.001 and were examined for association against each of the 4782 human serum protein measurements (see below).

### Protein measurements

Each protein has its own detection reagent selected from chemically modified DNA libraries, referred to as Slow Off-rate Modified Aptamers (SOMAmers)^55^. The design and quality control of the SOMApanel platform’s custom version to include proteins known or predicted to be present in the extracellular milieu have been described in detail elsewhere^10^. Briefly though, the aptamer-based platform measures 5034 protein analytes in a single serum sample, of which 4782 SOMAmers bind specifically to 4137 human proteins (some proteins are identified by more than one aptamer) and 250 SOMAmers that recognize non-human targets (47 non-human vertebrate proteins and 203 targeting human pathogens)^10^. Consistent target specificity across the platform was indicated by direct (through mass spectrometry) and/or indirect validation of the SOMAmers^10^. Both sample selection and sample processing for protein measurements were randomized, and all samples were run as a single set to prevent batch or time of processing biases.

### Statistical analysis

Prior to the analysis of the proteins measurements, we applied a Box-Cox transformation on all proteins to improve normality, symmetry and to maintain all protein variables on a similar scale^56^. In the association analysis, we obtained residuals after controlling for sex, age, potential population stratification using principal component (PCs) analysis^57^, and for all single-variant associations to serum proteins tested under an additive genetic model applying linear regression analysis (protein ~ SNP + age + sex + PC1 + PC2 + ….PC5). We report both variants to protein associations at P < 1×10^-6^ for suggestive evidence and Bonferroni correction for multiple comparisons by adjusting for the 54,469 variants and 4782 human protein analytes where single variant associations with P < 1.9×10^-10^ were considered study-wide significant (**Supplementary Table S1**). P-values corresponding to the estimated effect size and standard errors of the genotypes, were recalculated to increase accuracy. Independent genetic signals were found through a stepwise conditional and joint association analysis for each protein analyte separately with the GCTA-COJO software^58,59^. We conditioned on the current lead variant listed in **Supplementary Table S1**, defined as the variant with the lowest P-value, and then kept track of any new variants that were not in LD (the default GCTA-COJO option *r*^2^ < 0.9 for colinearity) with previously chosen lead variants and reported findings at P-value < 1×10^-6^ (**Supplementary Table S2**). In the joint model all conditionally significant SNPs for each protein analyte were combined in the regression model.

**Supplementary Table S3** summarizes, through use of VEP^24,25^, various pathogenicity prediction scores for all independent study-wide significant pQTLs in **Supplementary Table S2**, including the Likelihood Ratio Test (LRT)^60^, Variant Effect Scoring Tool (VEST)^61^, MutationAssessor^62^ and MutationTaster^63^. To test whether the percentage of secreted proteins among pQTLs is equal to the percentage of secreted proteins among non-pQTLs, 10,000 permutations were performed to obtain the empirical distribution of the χ^2^ test of equality of proportions. Our null and alternate hypotheses were:

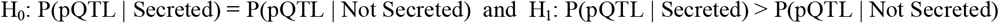

The test statistics calculated from our data was compared to the quantiles of this distribution to obtain P(Data|H_0_) (**Supplementary Fig. S1**).

We applied the “TwoSampleMR” R package^64^ to perform a two-sample MR analysis to test for causal associations between protein and outcome (protein-to-outcome). For different outcomes we used GWAS associations for LOAD in Europeans^32^, malignant melanoma in European individuals from the UK biobank data (UKB-b-12915)^65^ and T2D in Europeans^34^. Genetic variants (SNPs) associated with serum protein levels at a genome-wide significant threshold (P<5×10^-8^) identified in the AGES dataset and filtered to only include uncorrelated variants (*r*^2^<0.2) were used as instruments. The inverse variance weighted (IVW) method^66^ was used for the MR analysis, with P-values < 0.05 considered significant.

For the associations of individual proteins to different phenotypic measures we used linear or logistic regression or Cox proportional hazards regression, depending on the outcome being continuous, binary or a time to an event. Given consistency in terms of sample handling including time from blood draw to processing (between 9-11 am), same personnel handling all specimens and the ethnic homogeneity of the population we adjusted only for age and sex in all our regression analyses. All statistical analysis was performed using R version 3.6.0 (R Foundation for Statistical Computing, Vienna, Austria).

We compared our pQTL results to 19 previously published proteogenomic studies (**Supplementary Table 5**), including the protein GWAS in the INTERVAL study^11^, and our previously reported genetic analysis of 3,219 AGES cohort participants^10^. In the previous proteogenomic analysis of AGES participants, one *cis* variant was reported per protein using a locus-wide significance threshold, as well as *cis*-to-*trans* variants at a Bonferroni corrected significance threshold. Due to these differences in reporting criteria, we only considered the associations in previous AGES results that met the current study-wide P-value threshold. For all other studies we retained the pQTLs at the reported significance threshold. In addition, we performed a lookup of all independent pQTLs from the current study available in summary statistics from the INTERVAL study, considering them known if they reached a study-wide significance in their data. We calculated the LD structure between the reported significant variants for all studies, using 1000 Genomes v3 EUR samples, but using AGES data when comparing to previously reported AGES results. We considered variants in LD at r^2^>0.5 to represent the same signal across studies. Comparison was performed on protein level, by matching the reported Entrez gene symbol from each study.

## Acknowledgements

V.E. and Va.G. are supported by the Icelandic Research Fund (IRF grants 195761-051 and 184845-053). The Age, Gene/Environment Susceptibility-Reykjavik Study (AGES-RS) was supported by NIH contracts N01-AG-1-2100 and HHSN27120120022C, the NIA Intramural Research Program, Hjartavernd (the Icelandic Heart Association), and the Althingi (the Icelandic Parliament). M.A.K. was funded by Open Targets and by the Wellcome Trust Grant 206194.

## SUPPLEMENTARY MATERIAL

**Supplementary Fig. S1.**
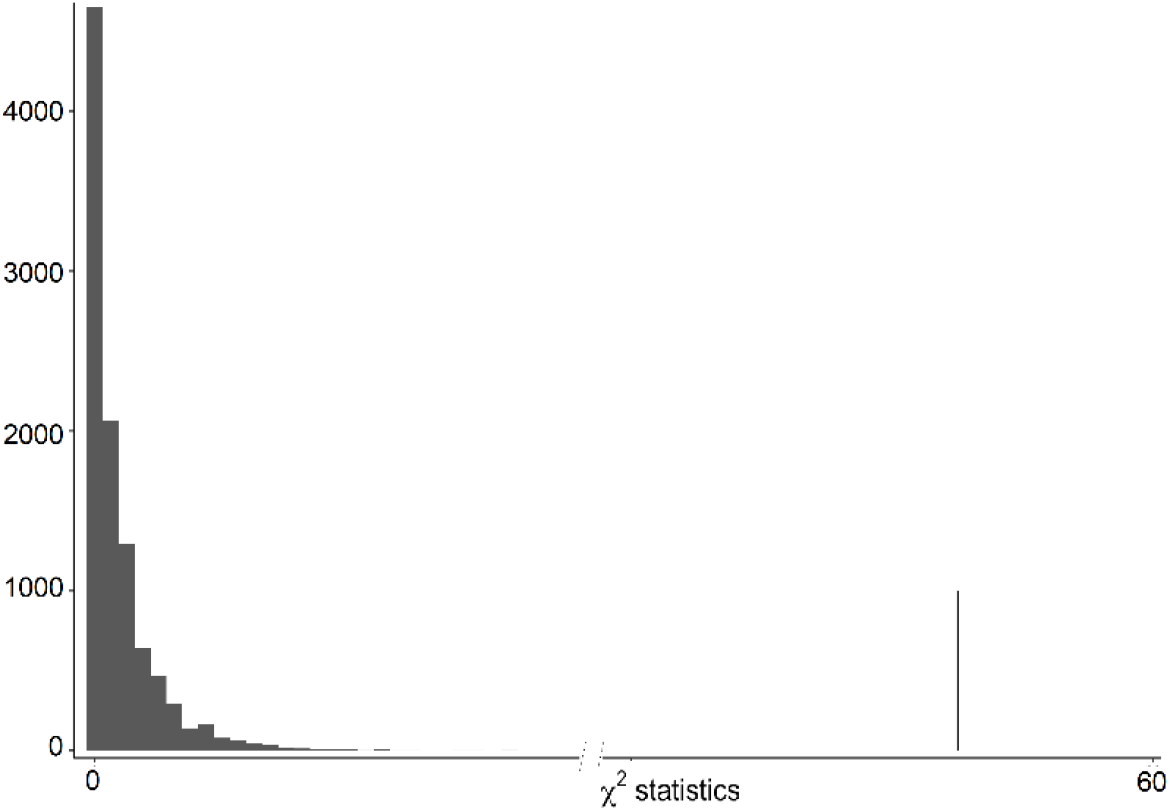
Empirical distribution of the test statistic as a histogram and the observed statistics calculated from our data as a vertical line. 10,000 permutations were performed to obtain the empirical distribution of the χ^2^ test of equality of proportions of pQTLs among secreted versus non-secreted proteins. Here, the test statistics calculated from our data to the quantiles of this distribution to obtain P(Data|H_0_) were compared. Of 10,000 permutations none gave a value greater than the observed statistic leading us to P-value = P(Data|H_0_) < 0.0001.

**Supplementary Fig. S2.**
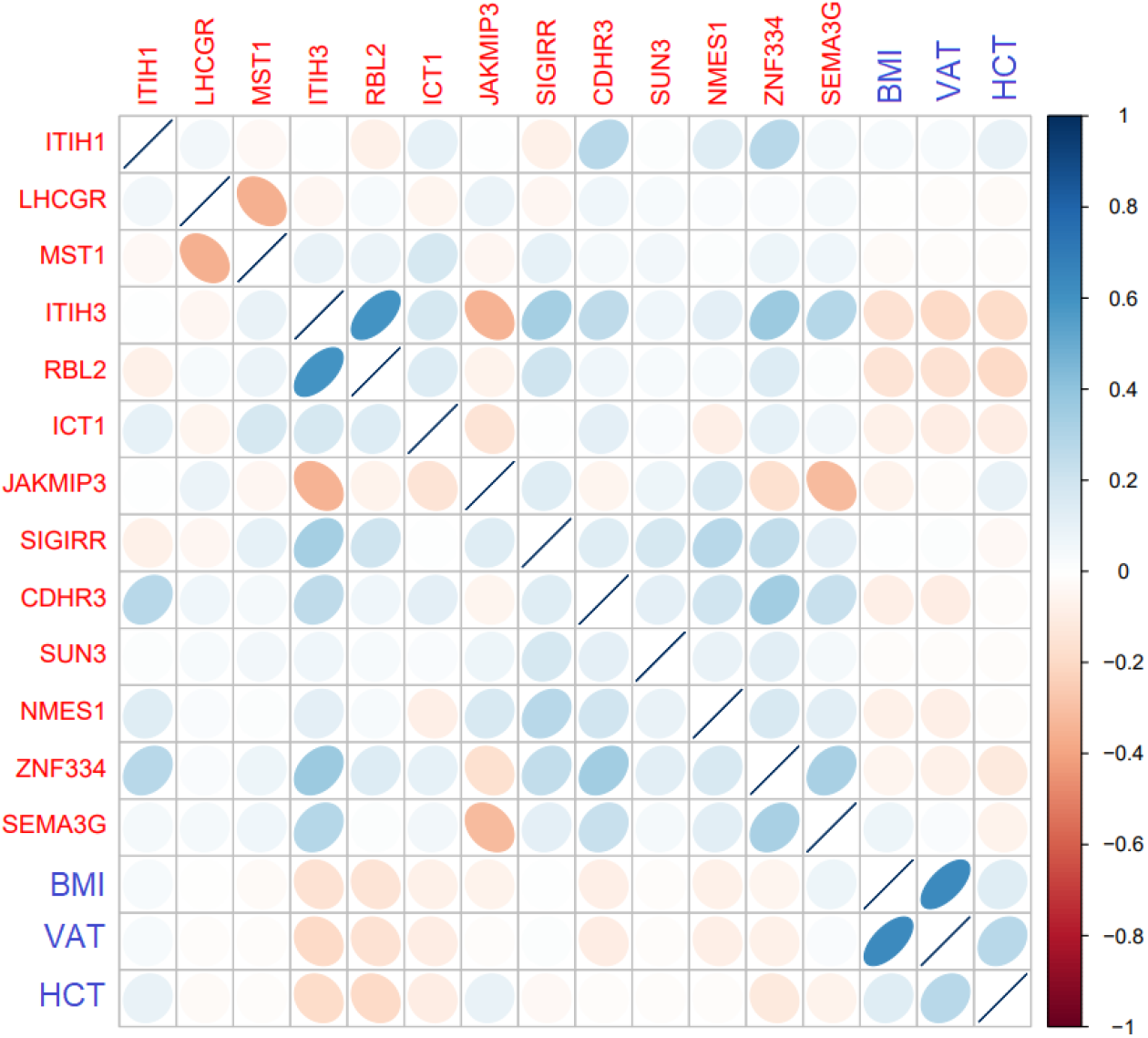
A Spearman rank correlation between all proteins as well as some quantitative traits including body mass index (BMI, kg/m2), visceral adipose tissue (VAT, measured *via* computed tomography) and hematocrit (HCT), that were associated with rs2251219.

**Supplementary Fig. S3.**
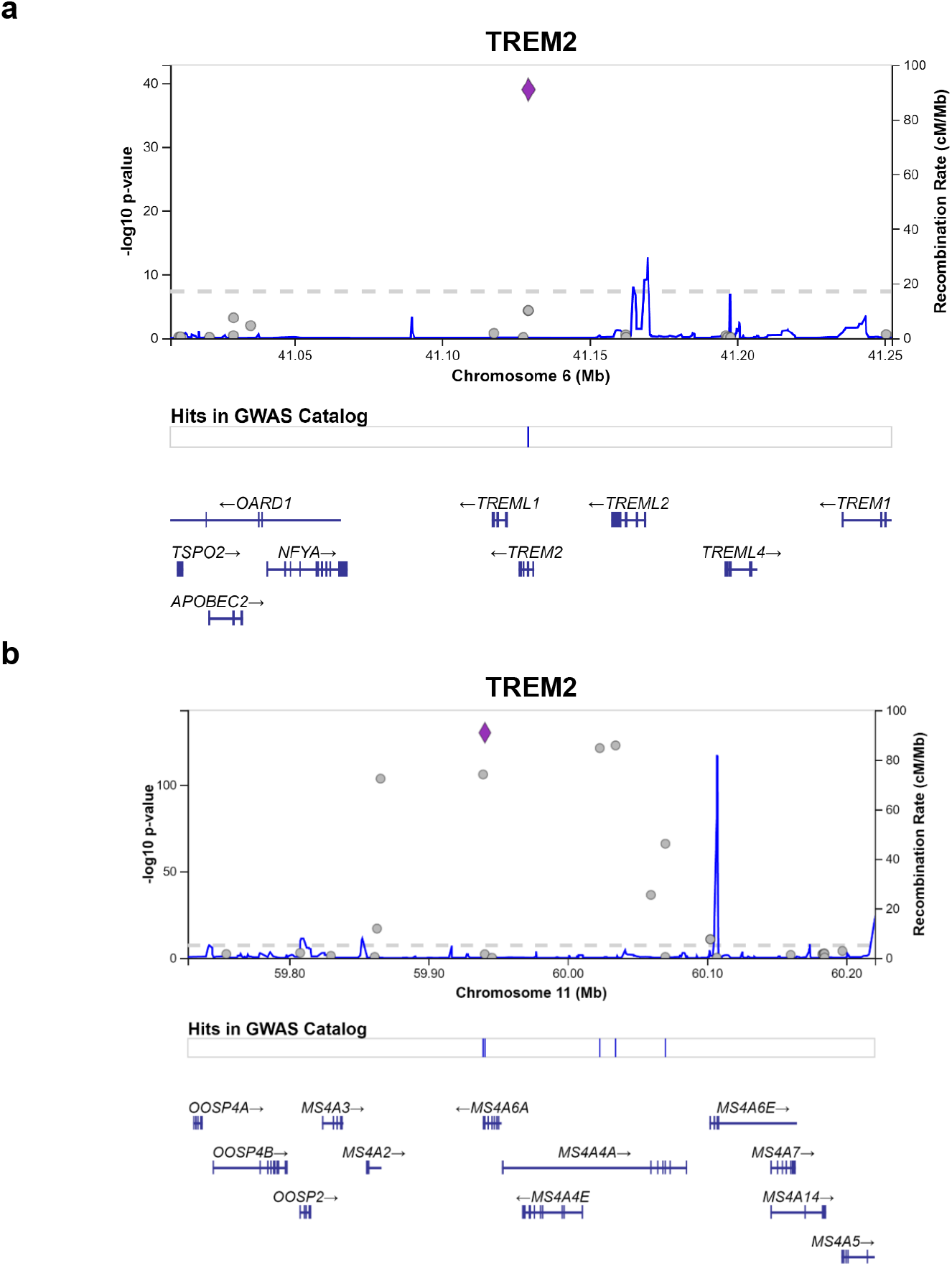
TREM2 regional plots (LocusZoom) based on exome array variants at chromosomes 6 and 11.

**Supplementary Fig. S4.**
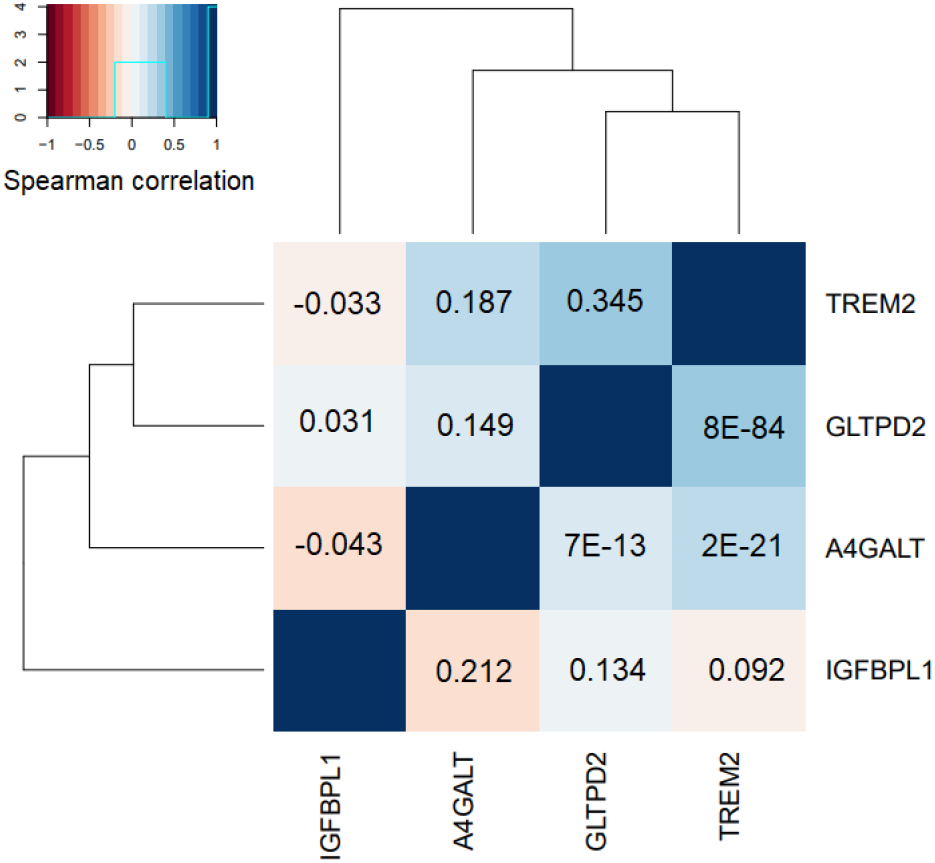
The graph shows the Spearman rank correlation between the four serum proteins affected by the two LOAD risk variants, rs75932628 and rs610932. The correlation matrix’s upper triangle depicts the beta-values, while the lower triangle highlights the P-values.

**Supplementary Fig. S5.**
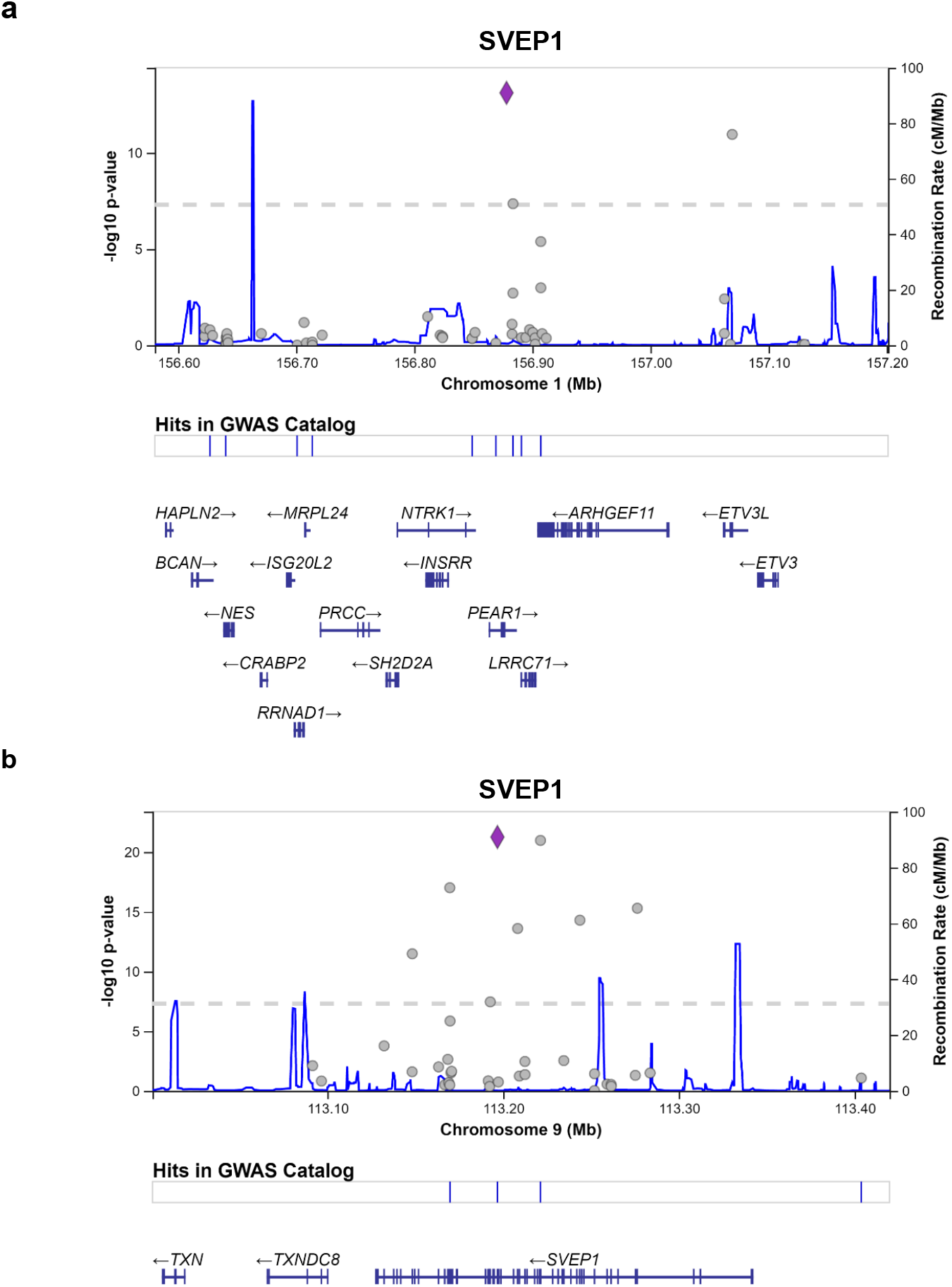
SVEP1 regional plots (LocusZoom) based on exome array variants at chromosomes 1 and 9.

## Notes

### Competing Interest Statement

J.R.L. and L.L.J. was and is, respectively, employees of and own stocks in Novartis. All other authors declare they have no competing interests.

## References

1 Buniello, A. et al. The NHGRI-EBI GWAS Catalog of published genome-wide association studies, targeted arrays and summary statistics 2019. Nucleic Acids Research 47, D1005–D1012, doi:10.1093/nar/gky1120 (2019).

2 Liu, D. J. et al. Exome-wide association study of plasma lipids in >300,000 individuals. Nature Genetics 49, 1758–1766, doi:10.1038/ng.3977 (2017).

3 Schadt, E. E. Molecular networks as sensors and drivers of common human diseases. Nature 461, 218–223, doi:10.1038/nature08454 (2009).

4 Zhang, B. et al. Integrated Systems Approach Identifies Genetic Nodes and Networks in Late-Onset Alzheimer’s Disease. Cell 153, 707–720, doi:10.1016/j.cell.2013.03.030 (2013).

5 Emilsson, V. et al. Genetics of gene expression and its effect on disease. Nature 452, 423–U422, doi:10.1038/nature06758 (2008).

6 Chen, Y. Q. et al. Variations in DNA elucidate molecular networks that cause disease. Nature 452, 429–435, doi:10.1038/nature06757 (2008).

7 Pires, D. E., Chen, J., Blundell, T. L. & Ascher, D. B. In silico functional dissection of saturation mutagenesis: Interpreting the relationship between phenotypes and changes in protein stability, interactions and activity. Sci Rep 6, 19848, doi:10.1038/srep19848 (2016).

8 Ho, J. E. et al. Common genetic variation at the IL1RL1 locus regulates IL-33/ST2 signaling. Journal of Clinical Investigation 123, 4208–4218, doi:10.1172/JCI67119 (2013).

9 Interleukin-6 Receptor Mendelian Randomisation Analysis, C. et al. The interleukin-6 receptor as a target for prevention of coronary heart disease: a mendelian randomisation analysis. Lancet 379, 1214–1224, doi:10.1016/s0140-6736(12)60110-x (2012).

10 Emilsson, V. et al. Co-regulatory networks of human serum proteins link genetics to disease. Science 361, 769–773, doi:10.1126/science.aaq1327 (2018).

11 Sun, B. B. et al. Genomic atlas of the human plasma proteome. Nature 558, 73–79, doi:10.1038/s41586-018-0175-2 (2018).

12 Mirauta, B. A. et al. Population-scale proteome variation in human induced pluripotent stem cells. Elife 9, doi:10.7554/eLife.57390 (2020).

13 Lamb, J. R., Jennings, L. L., Gudmundsdottir, V., Gudnason, V. & Emilsson, V. It’s in Our Blood: A Glimpse of Personalized Medicine. Trends Mol Med, doi:10.1016/j.molmed.2020.09.003 (2020).

14 Emilsson, V., Gudnason, V. & Jennings, L. L. Predicting health and life span with the deep plasma proteome. Nat Med 25, 1815–1816, doi:10.1038/s41591-019-0677-y (2019).

15 Lehallier, B. et al. Undulating changes in human plasma proteome profiles across the lifespan. Nat Med 25, 1843–1850, doi:10.1038/s41591-019-0673-2 (2019).

16 Williams, S. A. et al. Plasma protein patterns as comprehensive indicators of health. Nat Med 25, 1851–1857, doi:10.1038/s41591-019-0665-2 (2019).

17 Nakamura, A. et al. High performance plasma amyloid-beta biomarkers for Alzheimer’s disease. Nature 554, 249–254, doi:10.1038/nature25456 (2018).

18 Dodgson, S. E. There Will Be Blood Tests. Cell 173, 1–3, doi:10.1016/j.cell.2018.03.012 (2018).

19 Cohen, J. D. et al. Detection and localization of surgically resectable cancers with a multi-analyte blood test. Science 359, 926–930, doi:10.1126/science.aar3247 (2018).

20 Kristensen, S. L. et al. Prognostic Value of N-Terminal Pro-B-Type Natriuretic Peptide Levels in Heart Failure Patients With and Without Atrial Fibrillation. Circ Heart Fail 10, doi:10.1161/circheartfailure.117.004409 (2017).

21 Peloso, G. M. et al. Association of low-frequency and rare coding-sequence variants with blood lipids and coronary heart disease in 56,000 whites and blacks. American Journal of Human Genetics 94, 223–232, doi:10.1016/j.ajhg.2014.01.009 (2014).

22 Richards, A. L. et al. Exome arrays capture polygenic rare variant contributions to schizophrenia. Human Molecular Genetics 25, 1001–1007, doi:10.1093/hmg/ddv620 (2016).

23 Armengaud, J., Christie-Oleza, J. A., Clair, G., Malard, V. & Duport, C. Exoproteomics: exploring the world around biological systems. Expert Rev Proteomics 9, 561–575, doi:10.1586/epr.12.52 (2012).

24 McLaren, W. et al. The Ensembl Variant Effect Predictor. Genome Biol 17, 122, doi:10.1186/s13059-016-0974-4 (2016).

25 McLaren, W. et al. Deriving the consequences of genomic variants with the Ensembl API and SNP Effect Predictor. Bioinformatics 26, 2069–2070, doi:10.1093/bioinformatics/btq330 (2010).

26 Staley, J. R. et al. PhenoScanner: a database of human genotype-phenotype associations. Bioinformatics 32, 3207–3209, doi:10.1093/bioinformatics/btw373 (2016).

27 Jonsson, T. et al. Variant of TREM2 associated with the risk of Alzheimer’s disease. N Engl J Med 368, 107–116, doi:10.1056/NEJMoa1211103 (2013).

28 Guo, C. et al. IGFBPL1 Regulates Axon Growth through IGF-1-mediated Signaling Cascades. Sci Rep 8, 2054, doi:10.1038/s41598-018-20463-5 (2018).

29 Hollingworth, P. et al. Common variants at ABCA7, MS4A6A/MS4A4E, EPHA1, CD33 and CD2AP are associated with Alzheimer’s disease. Nature Genetics 43, 429–435, doi:10.1038/ng.803 (2011).

30 Suarez-Calvet, M. et al. sTREM2 cerebrospinal fluid levels are a potential biomarker for microglia activity in early-stage Alzheimer’s disease and associate with neuronal injury markers. EMBO Mol Med 8, 466–476, doi:10.15252/emmm.201506123 (2016).

31 Ewers, M. et al. Increased soluble TREM2 in cerebrospinal fluid is associated with reduced cognitive and clinical decline in Alzheimer’s disease. Sci Transl Med 11 (2019).

32 Kunkle, B. W. et al. Genetic meta-analysis of diagnosed Alzheimer’s disease identifies new risk loci and implicates Aβ, tau, immunity and lipid processing. Nat Genet 51, 414–430, doi:10.1038/s41588-019-0358-2 (2019).

33 Myocardial Infarction, G. et al. Coding Variation in ANGPTL4, LPL, and SVEP1 and the Risk of Coronary Disease. N Engl J Med 374, 1134–1144, doi:10.1056/NEJMoa1507652 (2016).

34 Mahajan, A. et al. Fine-mapping type 2 diabetes loci to single-variant resolution using high-density imputation and islet-specific epigenome maps. Nat Genet 50, 1505–1513, doi:10.1038/s41588-018-0241-6 (2018).

35 Nikpay, M. et al. A comprehensive 1,000 Genomes-based genome-wide association meta-analysis of coronary artery disease. Nat Genet 47, 1121–1130, doi:10.1038/ng.3396 (2015).

36 Evangelou, E. et al. Genetic analysis of over 1 million people identifies 535 new loci associated with blood pressure traits. Nat Genet 50, 1412–1425, doi:10.1038/s41588-018-0205-x (2018).

37 Huyghe, J. R. et al. Exome array analysis identifies new loci and low-frequency variants influencing insulin processing and secretion. Nature Genetics 45, 197–201, doi:10.1038/ng.2507 (2013).

38 Ransohoff, K. J. et al. Two-stage genome-wide association study identifies a novel susceptibility locus associated with melanoma. Oncotarget 8, 17586–17592, doi:10.18632/oncotarget.15230 (2017).

39 Lu, Y. et al. Large-Scale Genome-Wide Association Study of East Asians Identifies Loci Associated With Risk for Colorectal Cancer. Gastroenterology, doi:10.1053/j.gastro.2018.11.066 (2018).

40 Brown, K. M. et al. Common sequence variants on 20q11.22 confer melanoma susceptibility. Nature Genetics 40, 838–840, doi:10.1038/ng.163 (2008).

41 Blanchard, S. G. et al. Agouti antagonism of melanocortin binding and action in the B16F10 murine melanoma cell line. Biochemistry 34, 10406–10411 (1995).

42 Taylor, N. J. et al. Inherited variation at MC1R and ASIP and association with melanoma-specific survival. Int J Cancer 136, 2659–2667, doi:10.1002/ijc.29317 (2015).

43 Aguet, F. et al. The GTEx Consortium atlas of genetic regulatory effects across human tissues. bioRxiv, 787903, doi:10.1101/787903 (2019).

44 Wolf Horrell, E. M., Boulanger, M. C. & D’Orazio, J. A. Melanocortin 1 Receptor: Structure, Function, and Regulation. Front Genet 7, 95, doi:10.3389/fgene.2016.00095 (2016).

45 Zhang, B. et al. Large-scale genetic study in East Asians identifies six new loci associated with colorectal cancer risk. Nature Genetics 46, 533–542, doi:10.1038/ng.2985 (2014).

46 Calon, A. et al. Dependency of colorectal cancer on a TGF-beta-driven program in stromal cells for metastasis initiation. Cancer Cell 22, 571–584, doi:10.1016/j.ccr.2012.08.013 (2012).

47 Venkitachalam, S. et al. Biochemical and functional characterization of glycosylation-associated mutational landscapes in colon cancer. Sci Rep 6, 23642, doi:10.1038/srep23642 (2016).

48 Ishida, H. et al. A novel beta1,3-N-acetylglucosaminyltransferase (beta3Gn-T8), which synthesizes poly-N-acetyllactosamine, is dramatically upregulated in colon cancer. Febs Letters 579, 71–78, doi:10.1016/j.febslet.2004.11.037 (2005).

49 Solomon, T. et al. Identification of Common and Rare Genetic Variation Associated With Plasma Protein Levels Using Whole-Exome Sequencing and Mass Spectrometry. Circ Genom Precis Med 11, e002170, doi:10.1161/circgen.118.002170 (2018).

50 Smith, J. G. & Gerszten, R. E. Emerging Affinity-Based Proteomic Technologies for Large-Scale Plasma Profiling in Cardiovascular Disease. Circulation 135, 1651–1664, doi:10.1161/circulationaha.116.025446 (2017).

51 Zheng, J. et al. Phenome-wide Mendelian randomization mapping the influence of the plasma proteome on complex diseases. bioRxiv, 627398, doi:10.1101/627398 (2019).

52 Liu, B., Gloudemans, M. J., Rao, A. S., Ingelsson, E. & Montgomery, S. B. Abundant associations with gene expression complicate GWAS follow-up. Nature Genetics 51, 768–769, doi:10.1038/s41588-019-0404-0 (2019).

53 Harris, T. B. et al. Age, Gene/Environment Susceptibility-Reykjavik Study: multidisciplinary applied phenomics. Am J Epidemiol 165, 1076–1087, doi:10.1093/aje/kwk115 (2007).

54 Grove, M. L. et al. Best practices and joint calling of the HumanExome BeadChip: the CHARGE Consortium. PLoS One 8, e68095, doi:10.1371/journal.pone.0068095 (2013).

55 Candia, J. et al. Assessment of Variability in the SOMAscan Assay. Sci Rep 7, 14248, doi:10.1038/s41598-017-14755-5 (2017).

56 Max Kuhn, K. J. Applied Predictive Modeling. (Springer, 2013).

57 Price, A. L. et al. Principal components analysis corrects for stratification in genome-wide association studies. Nat Genet 38, 904–909, doi:10.1038/ng1847 (2006).

58 Yang, J. et al. Conditional and joint multiple-SNP analysis of GWAS summary statistics identifies additional variants influencing complex traits. Nat Genet 44, 369–375, s361-363, doi:10.1038/ng.2213 (2012).

59 Yang, J., Lee, S. H., Goddard, M. E. & Visscher, P. M. GCTA: a tool for genome-wide complex trait analysis. Am J Hum Genet 88, 76–82, doi:10.1016/j.ajhg.2010.11.011 (2011).

60 Chun, S. & Fay, J. C. Identification of deleterious mutations within three human genomes. Genome Res 19, 1553–1561, doi:10.1101/gr.092619.109 (2009).

61 Carter, H., Douville, C., Stenson, P. D., Cooper, D. N. & Karchin, R. Identifying Mendelian disease genes with the variant effect scoring tool. BMC Genomics 14 Suppl 3, S3, doi:10.1186/1471-2164-14-s3-s3 (2013).

62 Reva, B., Antipin, Y. & Sander, C. Predicting the functional impact of protein mutations: application to cancer genomics. Nucleic Acids Res 39, e118, doi:10.1093/nar/gkr407 (2011).

63 Schwarz, J. M., Rödelsperger, C., Schuelke, M. & Seelow, D. MutationTaster evaluates disease-causing potential of sequence alterations. Nat Methods 7, 575–576, doi:10.1038/nmeth0810-575 (2010).

64 Hemani, G. et al. The MR-Base platform supports systematic causal inference across the human phenome. Elife 7, doi:10.7554/eLife.34408 (2018).

65 Sudlow, C. et al. UK biobank: an open access resource for identifying the causes of a wide range of complex diseases of middle and old age. PLoS Med 12, e1001779, doi:10.1371/journal.pmed.1001779 (2015).

66 Burgess, S., Butterworth, A. & Thompson, S. G. Mendelian randomization analysis with multiple genetic variants using summarized data. Genet Epidemiol 37, 658–665, doi:10.1002/gepi.21758 (2013).

67 MacArthur, J. et al. The new NHGRI-EBI Catalog of published genome-wide association studies (GWAS Catalog). Nucleic Acids Research 45, D896–D901, doi:10.1093/nar/gkw1133 (2017).

